# Preoperative biliary stenting is associated with functional dysbiosis and impaired bile acid metabolism in pancreatic cancer - Impact of preoperative biliary stenting on gut microbiota

**DOI:** 10.64898/2026.05.19.719953

**Authors:** Marionna M. Cathomas, Eli Zamir, Marisa Isabell Keller, Toohina Gobin, Laila Jötten, Elias Gauer, Max Heckler, Bo Kong, Rogier Gaiser, Ingmar F. Rompen, Jonathan M. Harnoss, Sabine Schmidt, Franco Fortunato, Martin Loos, Eran Elinav, Peer Bork, Christoph W. Michalski, Thomas Hank

**Affiliations:** Department of General, Visceral and Transplantation Surgery, Heidelberg University Hospital, Heidelberg, Germany; Division of Microbiome and Cancer, German Cancer Research Center (DKFZ), Heidelberg, Germany; European Molecular Biology Laboratory (EMBL), Molecular Systems Biology Unit, Heidelberg, Germany; Department of Systems Immunology, Weizmann Institute of Science, Rehovot, Israel

**Keywords:** Pancreatic ductal adenocarcinoma, preoperative biliary stenting, intestine microbiome

## Abstract

Recent evidence suggests that the gut microbiome plays a role in the development and treatment response of pancreatic ductal adenocarcinoma (PDAC). However, the functional impact of tumor location and preoperative biliary stenting (PBS) on microbial composition and metabolism remains poorly understood. In this prospective study, preoperative stool specimens were collected from patients undergoing surgery for PDAC at Heidelberg University Hospital, Germany, between March 2020 and July 2021. Whole-genome shotgun metagenomic sequencing was performed to characterize microbial composition and functional pathways. A total of 63 preoperative stool samples were analyzed, including 40 patients with pancreatic head tumors (63.5%) and 23 with body/tail tumors (36.5%). Microbial community composition differed significantly according to tumor location (Bray–Curtis, p=0.005), with enrichment of Ruminococcus bromii in body/tail tumors. Among patients with pancreatic head tumors, PBS was associated with reduced alpha diversity (Shannon index, p=0.04), depletion of taxa including members of the Eubacteriales and Clostridiales orders as well as the genera Raoultella and Prevotella, and reduced abundance of selected genes involved in secondary bile acid metabolism. PBS was also associated with a higher rate of major postoperative complications according to Clavien–Dindo >3a (28.6% vs 3.8%; p=0.04). These findings suggest that biliary intervention may induce functional dysbiosis characterized by reduced microbial diversity and impaired bile acid metabolism, potentially disrupting host– microbiome crosstalk and contributing to adverse postoperative outcomes in pancreatic cancer.

## Introduction

The gut microbiome has emerged as an important modulator of host physiology and immune responses, with increasing evidence linking microbial alterations to tumor progression and adverse postoperative outcomes (1-3). Disruptions in microbial homeostasis can impair intestinal barrier function, promote bacterial translocation, and alter host–microbiome interactions, thereby contributing to systemic inflammation and disease progression (4). While these mechanisms have been extensively studied in colorectal surgery, their role in pancreatic ductal adenocarcinoma (PDAC) remains incompletely understood (5, 6).

In PDAC, tumor location is associated with distinct clinical and biological contexts (7). Tumors of the pancreatic head frequently present with obstructive jaundice, often requiring preoperative biliary stenting (PBS) (8). Beyond its clinical indication, PBS may influence the gut microbiome by altering bile flow and microbial metabolism, particularly pathways related to bile acid transformation. Such alterations may contribute to microbiome dysbiosis and impaired host–microbiome interactions (9). In contrast, tumors of the pancreatic body and tail typically develop without biliary obstruction, representing a different physiological environment.

Given the high postoperative morbidity and the dismal prognosis associated with PDAC, a better understanding of microbiome-associated mechanisms may identify modifiable targets to improve perioperative and, therefore, long-term outcomes (10). This study aimed to characterize taxonomic and functional differences in the gut microbiome of PDAC patients according to tumor location and to determine microbiome alterations associated with PBS.

## Methods

### Study design and population

This observational cohort study included 63 patients with PDAC located in the pancreatic head or body/tail, who underwent surgery between March 2020 and July 2021 at the Department of Surgery, Heidelberg University Hospital, Germany. Demographic and clinical data were retrieved from a prospectively maintained institutional database.

The study was approved by the Ethics Committee of Heidelberg University (S-141/2019). All patients provided informed consent.

### Follow-up and clinical variables

The primary endpoint was the assessment of differences in gut microbiota diversity between patients with PDAC located in the pancreatic head and those with body/tail tumors. A predefined subgroup analysis focused on patients with PDAC of the pancreatic head according to the presence of PBS, including an evaluation of microbial functional capacity with respect to secondary bile acid metabolism.

Patients received a single dose of antibiotics prior to PBS. In addition, patients undergoing PBS received an additional course of antibiotics on the day before surgery. By that time, however, the stool samples had already been collected.

Postoperative complications were defined as any medical or surgical condition deviating from the normal postoperative course and were graded according to the Clavien–Dindo classification (11). Complications graded ≥3a were considered major complications. All complications occurring within 30 days after hospital discharge, including reoperations and readmissions, were recorded.

### Collection and microbiome sequencing

Patients were recruited during outpatient consultation. After informed consent, stool samples were self-collected at home at least three days, but no more than 6 days, prior to surgery using a standardized collection kit and stored in Zymo DNA Shield solution. With the Zymo DNA shield solution, the samples could be stored at room temperature until processing. At the time of hospital admission, stool samples were collected for whole-genome sequencing.

### Sequencing data analysis

Raw sequencing reads were quality-filtered using fastp (version 0.23.4) to remove low-quality reads and reads shorter than 50 bp. Human contaminant reads were removed using Bowtie2 (version 2.3.5.1) with the hg38 reference genome and samtools (version 1.5). Remaining reads were rarefied to 25 million reads per sample using seqtk (version 1.4-r122). Taxonomic profiling was performed using Kraken (version 2.1.3), followed by species abundance re-estimation with Bracken (version 2.9), based on a Kraken database including human, mouse, and fungal GTDB references.

### Microbiome data analysis

Taxa with relative abundances below predefined thresholds were filtered as specified for each analysis, and remaining values below the threshold were set to the threshold value. Alpha diversity was calculated using the Shannon index as implemented in the shannon function of the skbio.diversity.alpha Python module. Beta diversity was assessed after abundance filtering (threshold 0.001) using Bray-Curtis dissimilarity and visualized by principal coordinate analysis (PCoA) with the skbio.stats.distance and skbio.stats.ordination modules.

Differential abundance analysis was performed after abundance filtering (threshold 0.01) using the Mann– Whitney U test. Species with a p-value <0.05 and a fold change >1.3 were considered significant. The differential abundance analysis was applied on data which was normalized to a sum of 1 per sample, without further subjecting it to centered log-ratio (CLR) transformation. The testing was based on a non-parametric test (Mann-Whitney U rank test). For subgroup analyses, microbial functional potential was assessed using Kyoto Encyclopedia of Genes and Genomes (KEGG) pathway analysis. Secondary bile acid metabolism was evaluated using KEGG pathway ko00121, focusing on K01442 (bile salt hydrolase), K07007 (baiN), and K15969 (baiB).

### Statistical analysis

Statistical analyses were performed using SPSS Statistics (version 30). Continuous variables were analyzed using Student’s t-test or Mann–Whitney U test, as appropriate, and are presented as mean or median values. Categorical variables were compared using the chi-square test or Fisher’s exact test, as indicated. Binary logistic regression analysis was used to identify independent predictors, with results expressed as odds ratios. A two-sided p value <0.05 was considered statistically significant.

Analysis of variance (ANOVA) was applied to assess differences in Shannon diversity. Differences in alpha diversity were additionally evaluated using the Mann–Whitney U test as implemented in the scipy.stats module. Differences in beta diversity were assessed using permutational multivariate analysis of variance (PERMANOVA) implemented in skbio.stats.distance. Functional analysis was performed on KEGG-PATHWAY and keg-orthologous genes (KO) (prevalence >= 0.05, abundance >= 1e-5) using wilcoxon rank sum test with subsequent false discovery rate (fdr) correction for multiple testing.

## Results

### Patient cohort and characteristics

A total of 63 patients with PDAC were included, comprising tumors located in the pancreatic head (n=40) and body/tail (n=23). Baseline demographic and tumor-related characteristics were comparable between groups (**Tables 1–2**). Patients with pancreatic head tumors underwent more complex surgical procedures and had longer operative times (p=0.024). Major postoperative complications >3a according to Clavien-Dindo classification occurred more frequently in patients with pancreatic head tumors (17.5% vs 0%; p=0.031), with a corresponding increase in 90-day mortality (7.5% vs. 0; p=0.546).

**Table 1:** Patients characteristics of the cohort. BMI, body mass index in kg/m, ASA, Grading in American Society of Anesthesiologists; PPI, proton pump inhibitor

|  | All patients<br>(n=63) | PDAC head<br>(n=40) | PDAC body/tail<br>(n=23) | p-value |
| --- | --- | --- | --- | --- |
| **Age (median; IQR)** | 64 (43-80) | 64 (48-80) | 66 (43-77) | 0.721 |
| **Gender (male)** | 43 (68.3%) | 28 (70.0%) | 15 (65.2%) | 0.695 |
| **Weight (median; IQR)** | 78 (48-105) | 78.1 (48-105) | 77.3 (55-103.3) | 0.618 |
| **BMI (median; IQR)** | 25.3 (17.6-37.9) | 25.4 (17.6-32.5) | 25.1 (18.8-37.9) | 0.618 |
| **ASA** |  |  |  |  |
| 2 | 38 (60.3%) | 23 (57.5%) | 15 (65.2%) | 0.547 |
| 3 | 25 (39.7%) | 17 (42.5%) | 8 (34.8%) |  |
| **Diabetes** |  |  |  |  |
| No | 44 (69.8%) | 27 (67.5%) | 17 (73.9%) | 0.593 |
| Yes | 13 (31.7%) | 13 (32.5%) | 19 (26.1%) |  |
| **Metformin** |  |  |  |  |
| No | 52 (82.5%) | 34 (85.0%) | 18 (78.3%) | 0.498 |
| Yes | 11 (17.5%) | 6 (15.0%) | 5 (21.7%) |  |
| **PPI** |  |  |  |  |
| No | 44 (69.8%) | 26 (65.0%) | 18 (78.3%) | 0.270 |
| Yes | 19 (30.2%) | 14 (35.0%) | 5 (21.7%) |  |

**Table 2:**
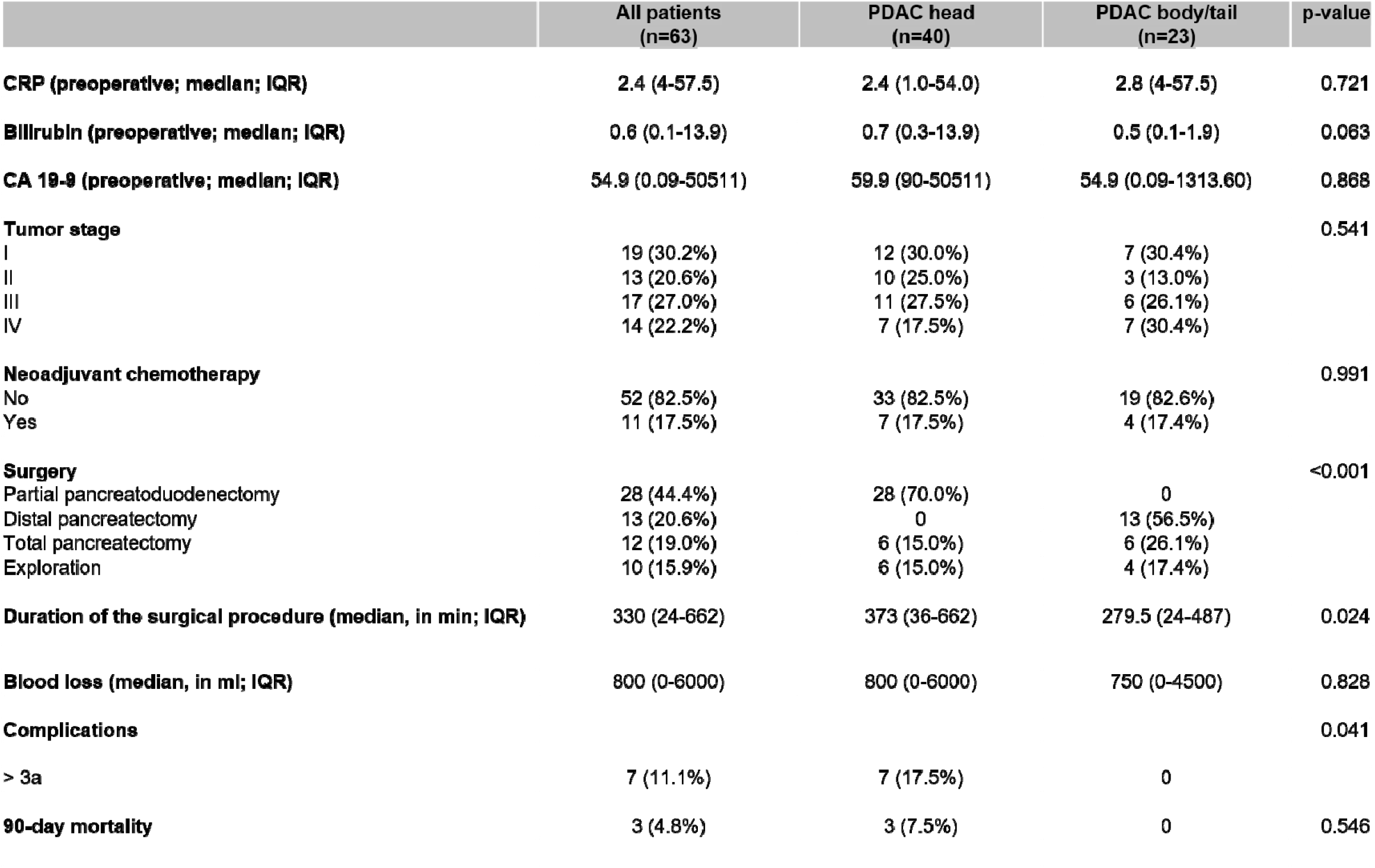
Surgical and clinical outcomes. CA 19-9, carbohydrate antigen 19-9; CRP, C-reactive protein * According to Clavien-Dindo classification within 30 days after discharge

### Microbiome differences by tumor location

Microbial alpha diversity (Shannon index) of stool samples did not differ between pancreatic tumor locations (p=0.21; **Figure 1A**). In contrast, microbial community composition differed significantly between groups, as assessed by Bray–Curtis dissimilarity (p=0.005; **Figure 1B**). Differential abundance analysis identified an enrichment of *Ruminococcus bromii* in patients with pancreatic body/tail tumors (**Figure 2**).

**Figure 1:**
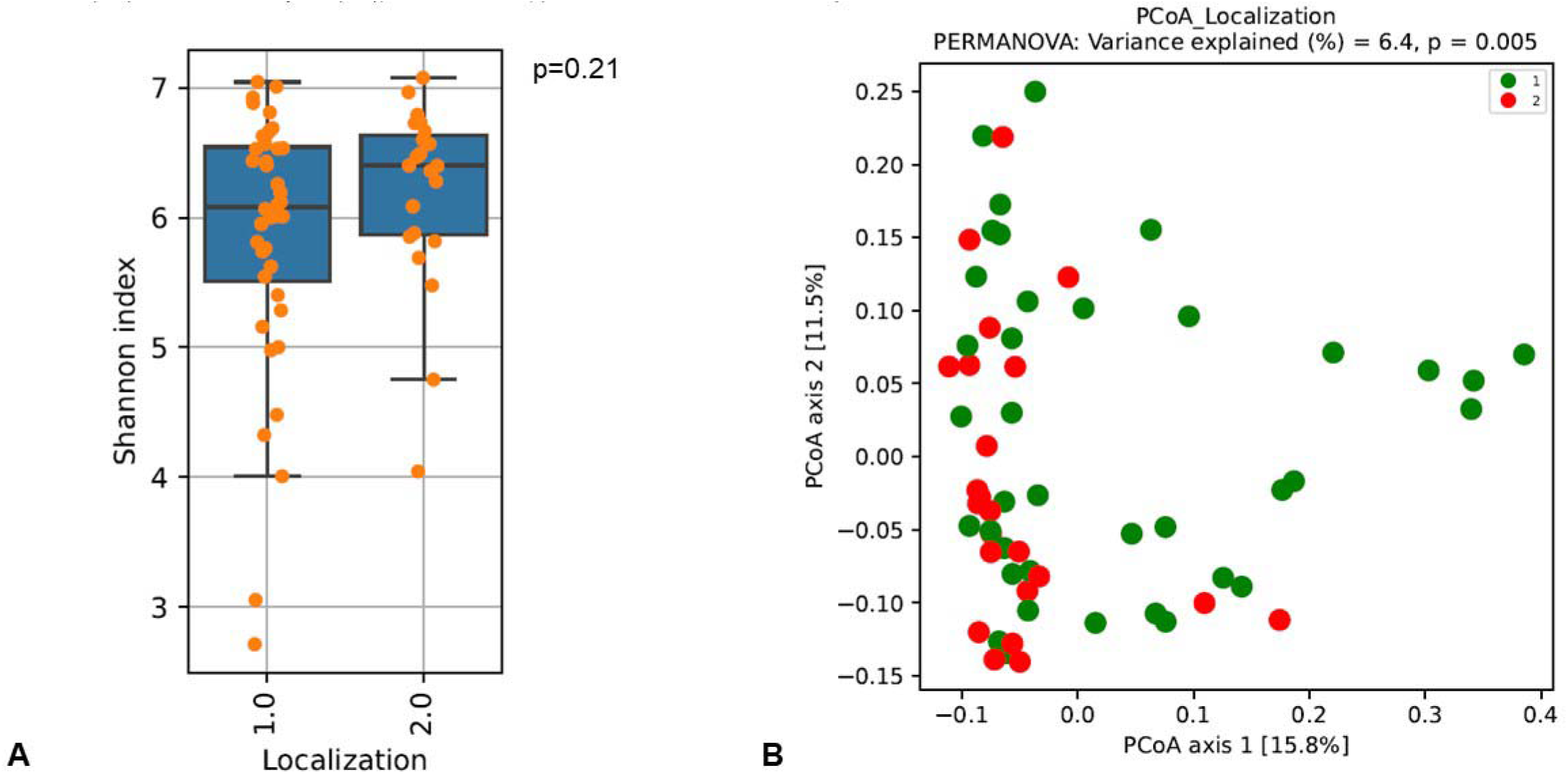
Alpha diversity (A) measured by Shannon index and beta diversity (B) analyzed by principal coordinate analysis (PCoA) based on species-level Bray-Curtis dissimilarly. PDAChead (1.0) and PDAC body/tail (2.0); PERMANOVA, permutational multivariate analysis of variance.

**Figure 2:**
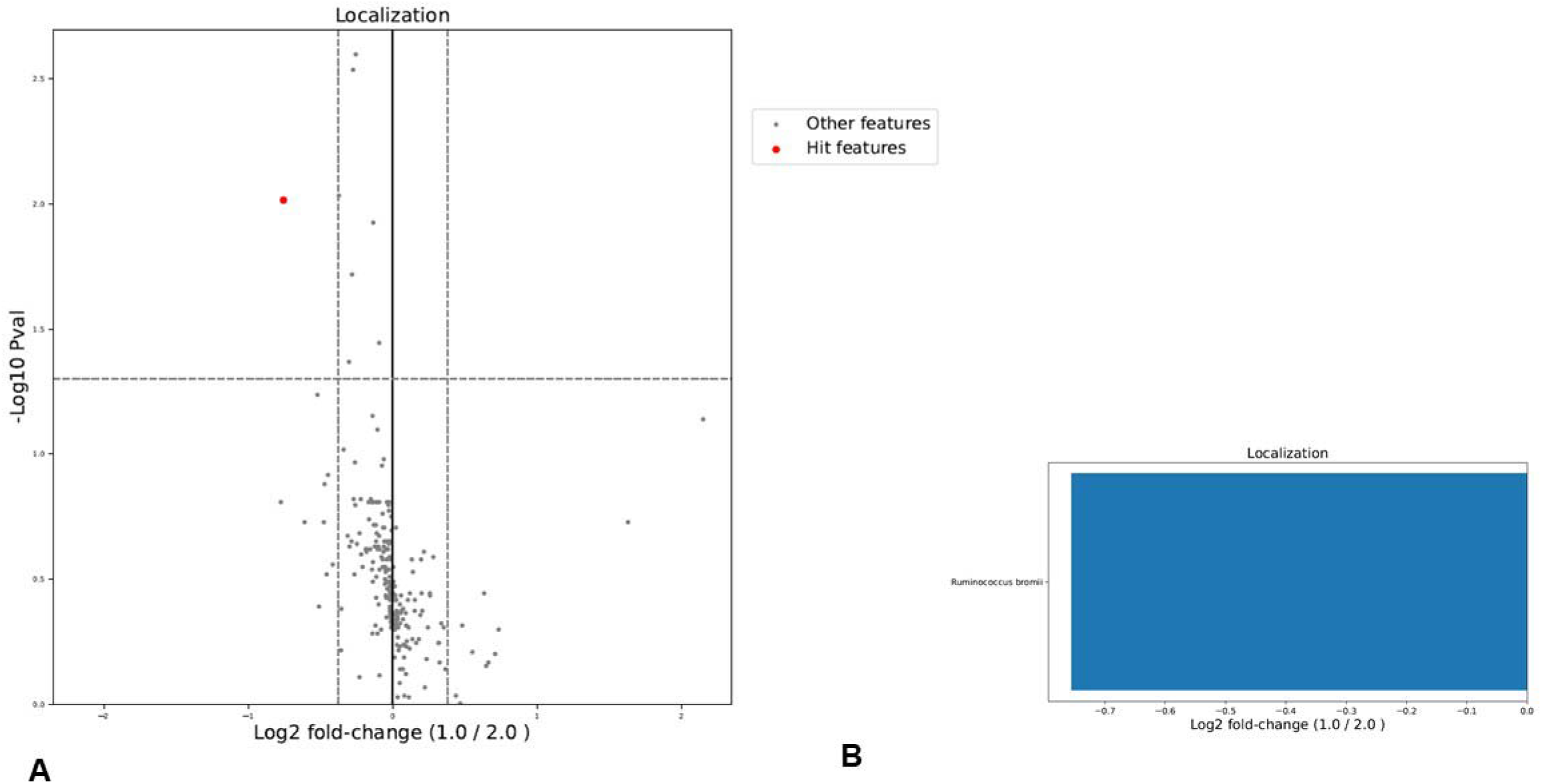
A) Relative abundance in PDAC head (1.0) and PDAC body/tail (2.0). B) Taxa with significance higher abundance in patients with PDAC body/tail.

### Clinical correlations of tumor locations and PBS

Among patients with PDAC located in the head, 14 (35.0%) underwent PBS. Baseline characteristics and surgical variables were comparable between patients with and without PBS (**Suppl. Tables 2-3)**. Further, all known influencing factors as metformin or proton pump inhibitor (PPI) were comparable among the two groups. The median interval between PBS and surgery was 29 days (IQR: 18.0-66.5). Patients with PBS experienced a higher rate of major postoperative complications compared with those without PBS (28.6% vs 3.8%; p=0.04;). Intraoperative bile cultures, available in 67.5% of patients, were more frequently positive in the PBS group (85.7% vs 42.3%; p<0.001), with predominantly polymicrobial growth (**Table 3**).

**Table 3:**
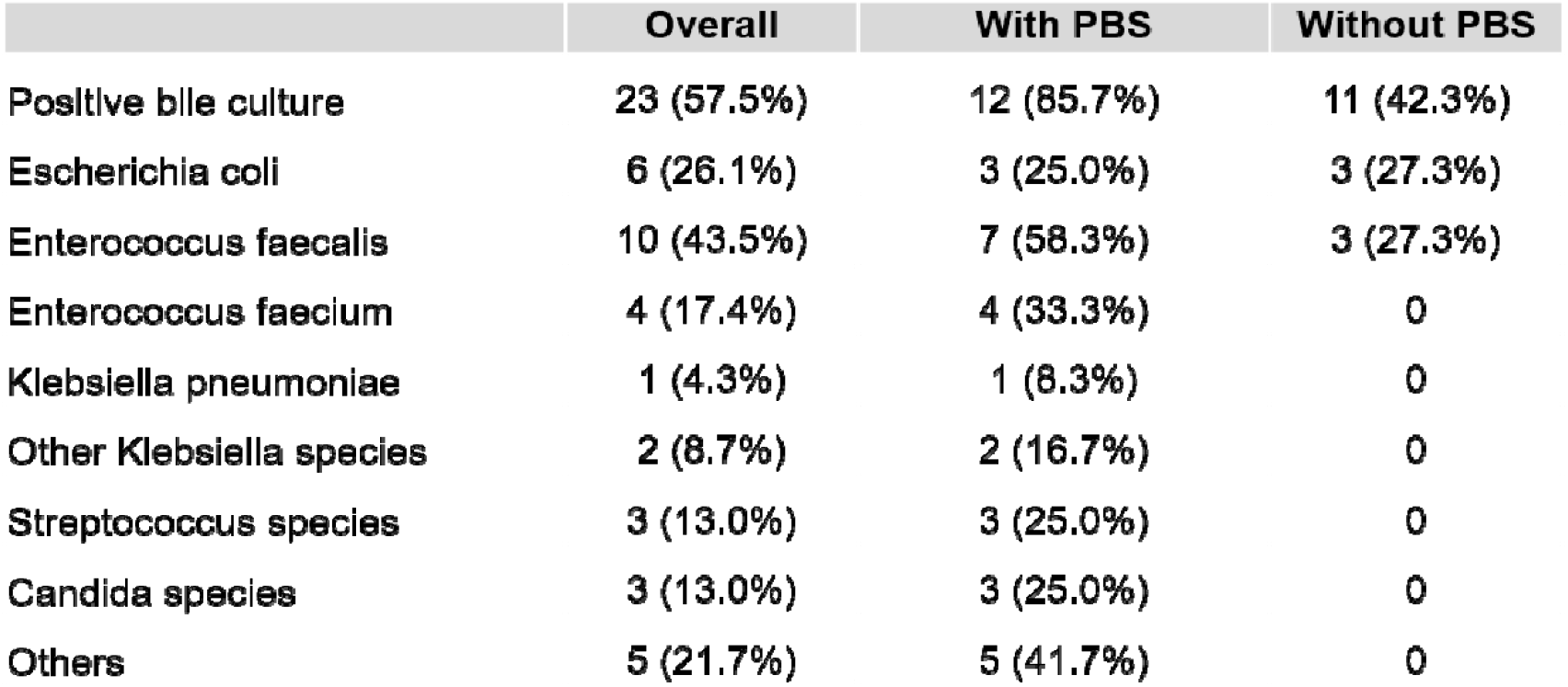
Microbiological analysis of positive bile culture in patient with and without preoperative biliary stent (PBS)

### Microbiome alterations associated with PBS

PBS was associated with reduced alpha diversity (p=0.04; **Figure 3A**), whereas no significant difference in beta diversity was observed (p=0.138; **Figure 3B**). Taxonomic analysis demonstrated a relative depletion of taxa commonly associated with gut homeostasis, including *Eubacteriales* and *Clostridiales*, in patients with PBS (**Figure 4**). Reduced alpha diversity was associated with the occurrence of major postoperative complications in multivariable analysis (p=0.04; **Suppl. Table 1**).

**Figure 3:**
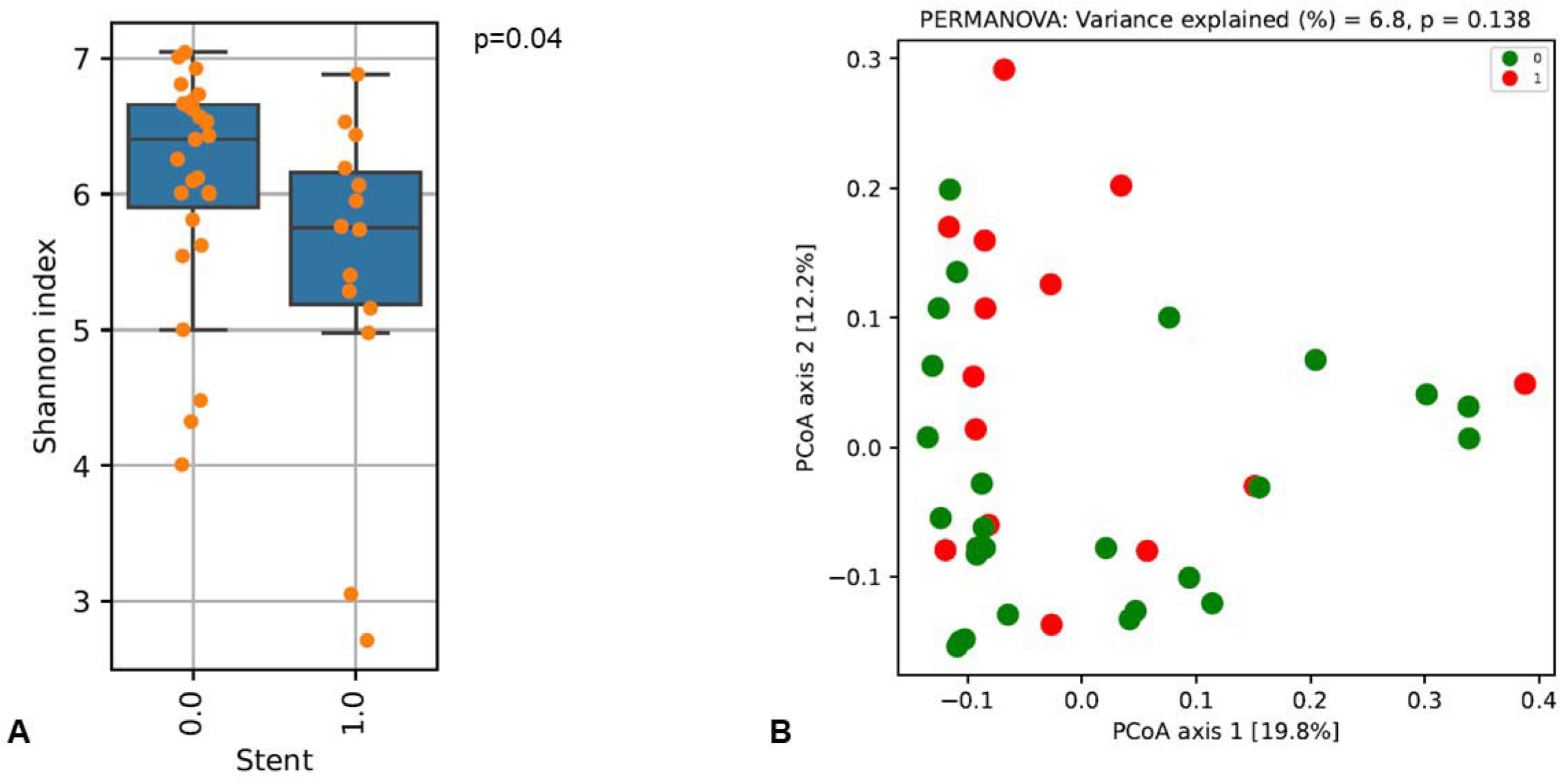
Alpha diversity (A) measured by Shannon index and beta diversity (B) analyzed by principal coordinate analysis (PCoA) based on species-level Bray-Curtis dissimilarity. PDAC head without (0.0) and with (1.0) preoperative biliary stent; PERMANOVA, permutational multivariate analysis of variance.

**Figure 4:**
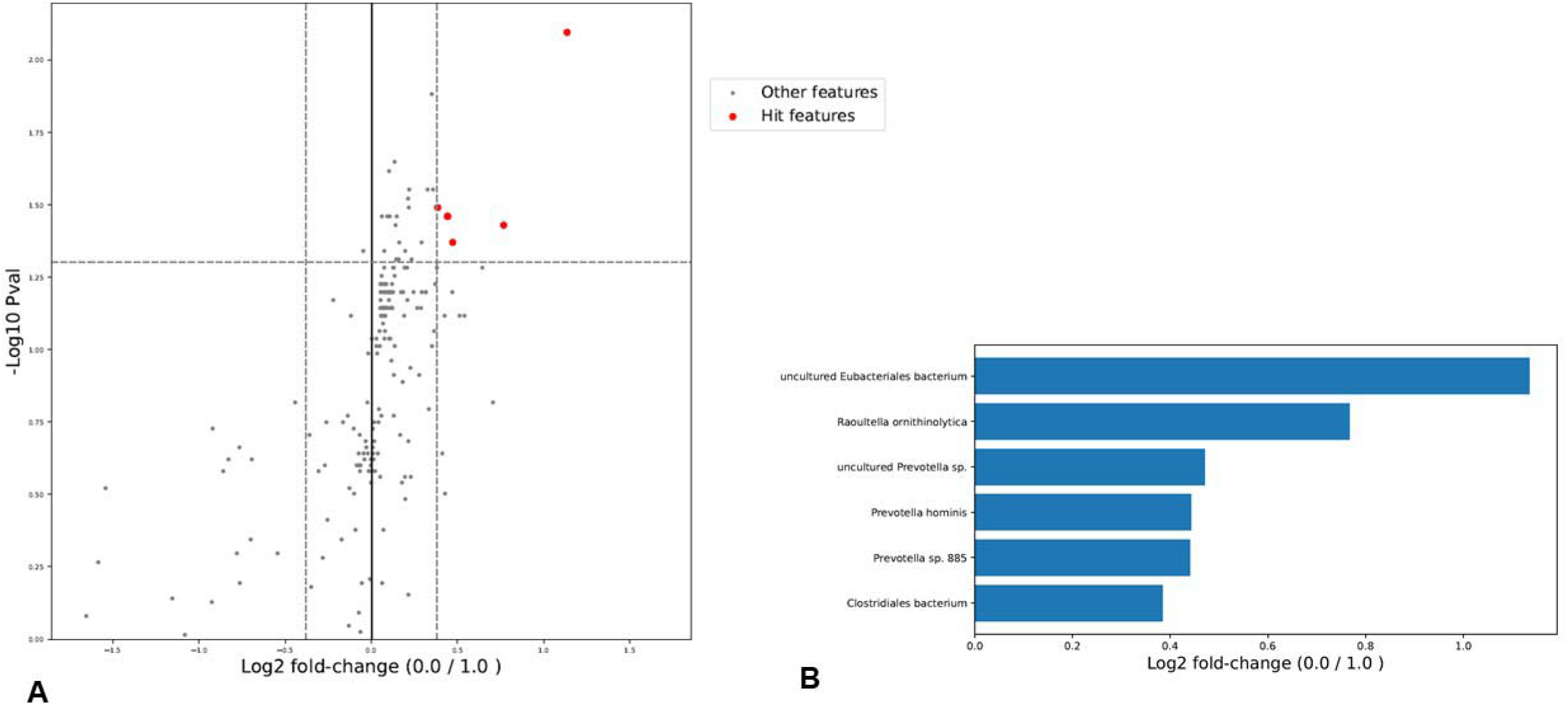
A) Relative abundance in patients without (0.0) and with (1.0) preoperative biliary stent. B) Taxa with significance higher abundance in patients without preoperative biliary stent.

### Functional microbiome profiling

To further explore the biological relevance of these microbial alterations, functional pathway analysis was performed on 4443 KEGG orthologous genes. We found no significantly changing genes after correcting for multiple testing. Because PBS could likely have an impact on the bile acid compositions in the large intestine, we especially examined the bile acid-related KOs and functional pathways abundances. Functional pathway analysis revealed no change in the secondary microbial bile acid metabolism pathway (map00121) in patients with PBS (**Figure 5A**). Also individual KO within the pathways, the baiN-associated gene (K07007), the bile salt hydrolase (K01442) or the baiB gene (K15969) show no difference (**Figure 5B**). These findings indicate that the secondary microbial bile acid metabolism stays stable in patients undergoing PBS.

**Figure 5:**
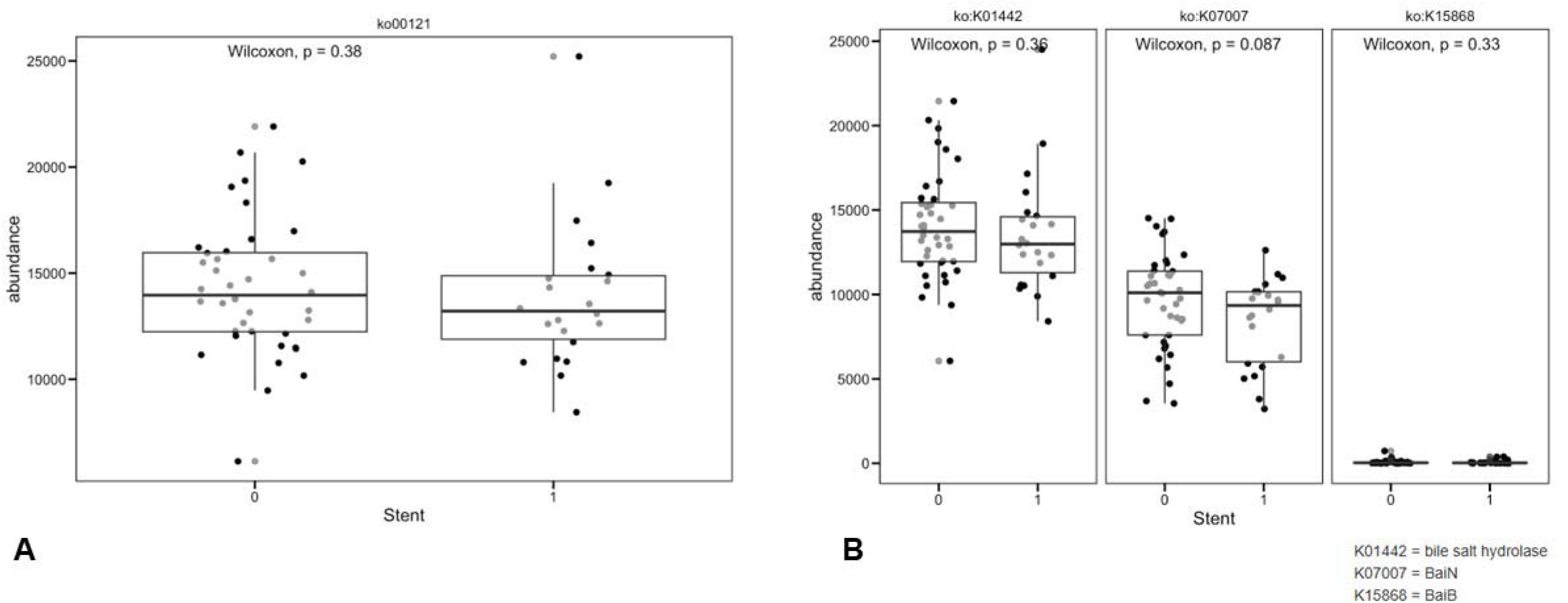
Secondary bile acid metabolism using Kyoto Encyclopedia of Genes and Genomes (KEEG) pathway (A). Functional activity was measured by K01442 (bile salt hydrolase), K07007 (baiN) and K15969 (baiB) (B). Assessment was performed in patients with and without preoperative biliary stent (PBS).

## Discussion

In this study, we demonstrate that tumor location and PBS are associated with distinct alterations of the preoperative gut microbiome in PDAC. While tumors located in the pancreatic body and tail were characterized by differences in microbial community composition, PBS in patients with pancreatic head tumors was associated with reduced within-patient microbial diversity and but no impaired microbial bile acid metabolism. These findings suggest that tumor-related and iatrogenic factors are linked to distinct microbiome alterations with potential clinical relevance.

Patients with PDAC located in the pancreatic body and tail showed significantly different beta diversity compared with those with tumors in the pancreatic head, while alpha diversity remained comparable. This indicates that tumor location primarily affects microbial community composition rather than overall microbial richness. The observed enrichment of *Ruminococcus bromii* in body/tail tumors is noteworthy, as this species plays a central role in the degradation of resistant starch and supports the production of short-chain fatty acids such as butyrate (12). Through these mechanisms, *Ruminococcus bromii* has been linked to microbial stability, maintenance of epithelial barrier integrity and modulation of immune responses. Accordingly, the presence of such taxa may reflect a microbial profile more consistent with gut homeostasis. (13).

These location-dependent microbial differences are biologically plausible. Pancreatic head tumors typically arise in a biliary-influenced environment and are frequently associated with cholestasis and endoscopic interventions, whereas body and tail tumors develop largely independent of direct biliary exposure. These findings suggest a more gut-derived microbial profile in body/tail tumors, while head tumors appear to be more strongly influenced by biliary conditions (8, 14-17). While surgical complexity likely explains the higher complication rates observed in patients with pancreatic head tumors, microbiome-associated factors may additionally contribute to postoperative vulnerability (18).

In contrast to tumor location, PBS was associated with a reduction in alpha diversity, indicating a loss of microbial richness. Notably, beta diversity remained comparable, suggesting a global depletion of microbial diversity rather than a distinct microbial community shift. This pattern may be consistent with a microbiome profile suggestive of dysbiosis, potentially mediated by retrograde bacterial translocation, biofilm formation or prolonged exposure of the biliary system to intestinal microbes (19). These alterations suggest a disruption of host–microbiome crosstalk driven by biliary intervention (14).

Importantly, PBS was not associated with functional changes of the gut microbiome, particularly the secondary bile acid metabolism pathway and gene abundances stayed consistent (17). Secondary bile acids play a key role in maintaining intestinal homeostasis and regulating inflammatory responses (20, 21). Their consistency may therefore represent a compensation between biliary intervention and microbiome alterations (22, 23). These findings suggested preoperative biliary stenting may represent a potential iatrogenic driver of functional dysbiosis.

These findings are in line with previously published data demonstrating that PBS are associated with altered biliary and intestinal microbiota and higher rates of major complications (24). While causality cannot be established, the association between reduced microbial diversity, consistent bile acid metabolism, and adverse clinical outcomes suggests that microbiome changes may reflect a vulnerable perioperative state. Notably, bile culture positivity alone was not associated with complications, supporting the concept that global microbiome alterations may be more informative than individual pathogens (25, 26).

From a clinical perspective, both tumor location and PBS are identifiable preoperatively, but only PBS represents a potentially modifiable factor (27). Current guidelines already recommend restrictive use of preoperative biliary drainage; our findings provide additional biological context support for this approach. In the future, microbiome-based risk stratification and targeted perioperative interventions may represent promising strategies, although prospective studies are required. (8, 28).

This study has several limitations. The cohort size is limited, particularly for subgroup analyses and the observational design precludes causal inference. Stool-based microbiome profiling does not capture site-specific microbial dynamics. Dietary intake and other environmental factors influencing the microbiome could not be fully controlled. In addition, functional predictions based on metagenomic data require validation by metabolomic analyses. Moreover, the absence of non-PDAC controls limits conclusions regarding disease-specific microbial signatures. Accordingly, our findings should be interpreted as differences within PDAC according to tumor location and PBS.

In conclusion, tumor location and PBS are associated with distinct alterations of the gut microbiome in PDAC. While tumor location primarily influences microbial composition, biliary stenting is linked to reduced microbial diversity. These findings support the possible interaction between clinical factors and the gut microbiome and suggest that microbiome alterations induced by biliary intervention may represent a modifiable biological pathway linking clinical practice to postoperative risk.

## Acknowledgements

We acknowledge the EMBL Genomics Core Facility for support in metagenomic sequencing.

## Qualify for (co-) authorship

a. Conception and design, or analysis and interpretation of data; MC, EZ, MIK, TH
b. Drafting the article or revising it critically for important intellectual content; MC, EZ, MIK, RG, FF, EE, PB, TH
c. Final approval of the version to be published. MC, LJ, EG, MH, BK, IFR, JMH, SS, FF, ML, EE, CWM, TH

## Notes

### Competing Interest Statement

The authors have declared no competing interest.

