## Supplementary material for "Preoperative biliary stenting is associated with functional dysbiosis and impaired bile acid metabolism in pancreatic cancer - Impact of preoperative biliary stenting on gut microbiota": Suppl. Tables 1-3

**Supplementary data, table 1: Multivariable analysis to analyse independent factor for low alpha diversity**

Positive predictive factors was preoperative biliary stent.

CI, confidence interval; BMI, body mass indey; CA 19-9, carbohydrate antigen 19-9; CEA, carcinoembyonic antigen

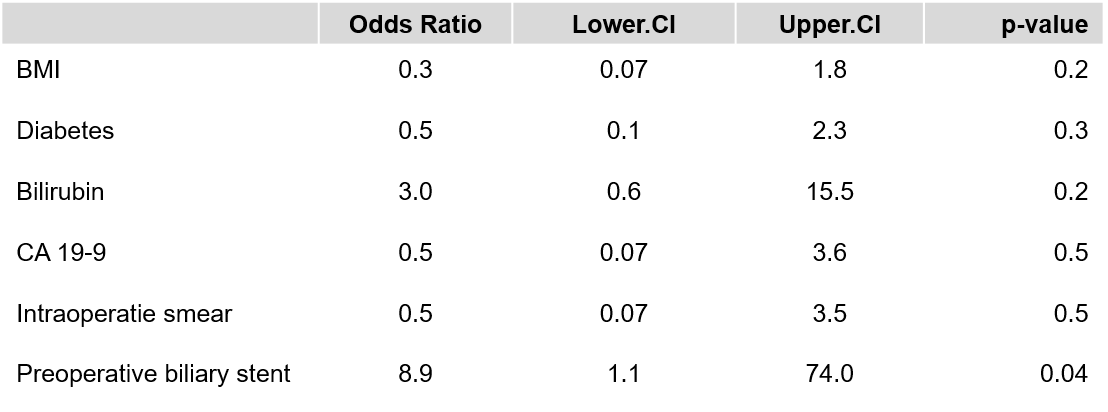

**Supplementary data, Table 2: Patients characteristics of the cohort**

PBS, preoperative biliary stent; BMI, body mass index in kg/m^2^, ASA, Grading in American Society of Anesthesiologists; PPI, proton pump inhibitor

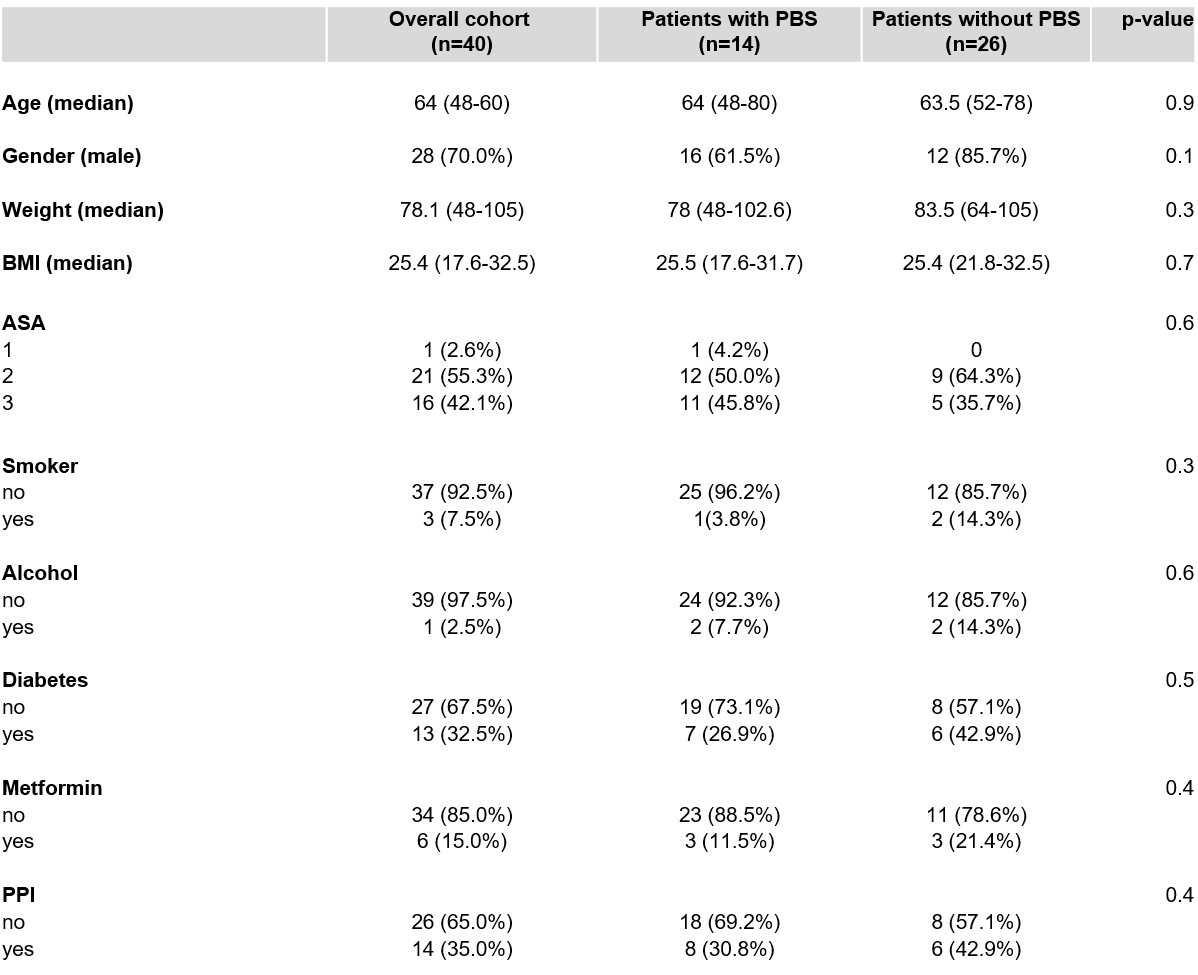

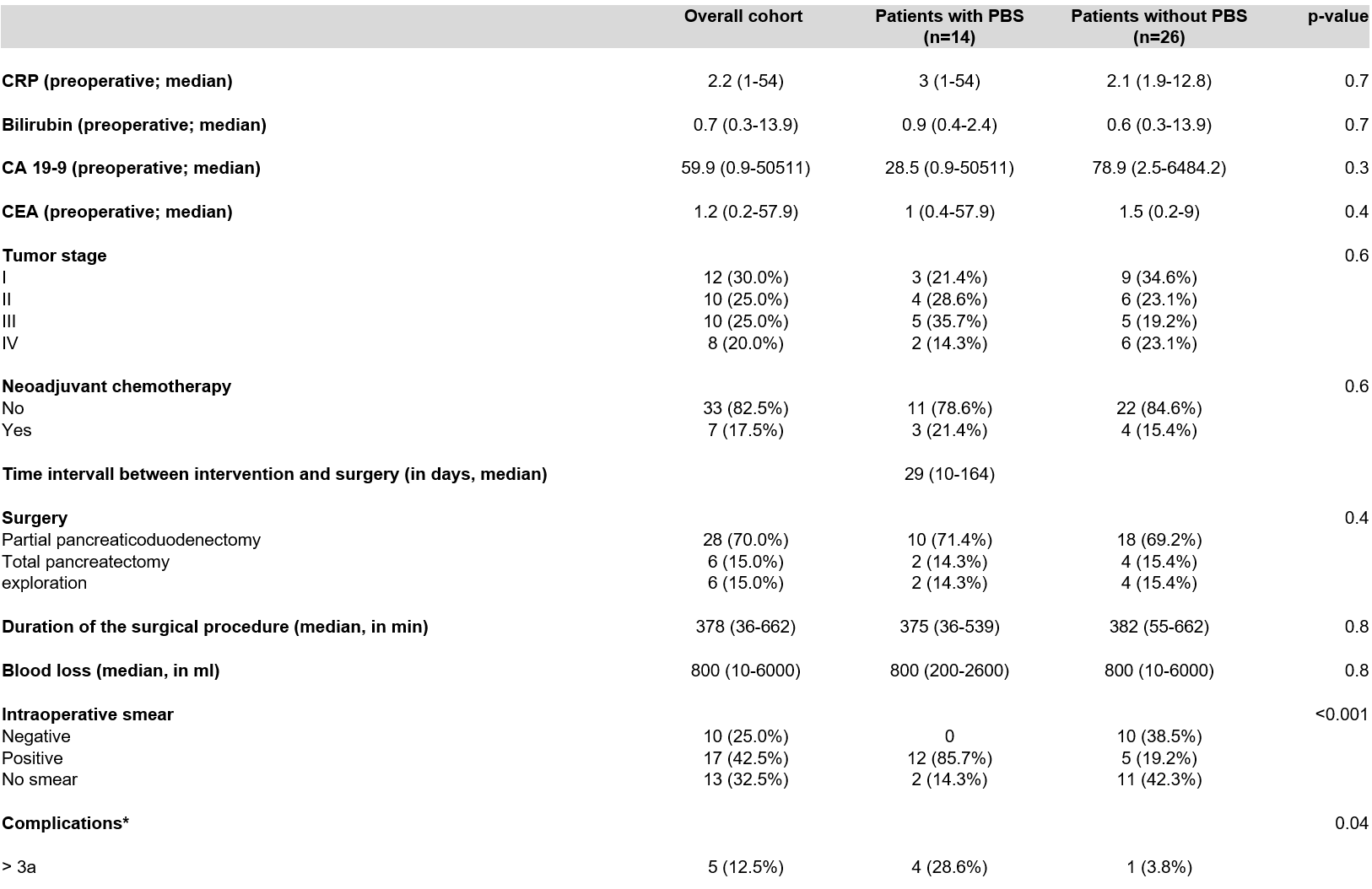

**Supplementary data, Table 3: Surgical and clinical outcomes**

PBS, preoperative biliary stent; CA 19-9, carbohydrate antigen 19-9; CEA, carcinoembyonic antigen* According to Clavien-Dindo classification within 30 days after discharge
